## Supplemental Figures for "Structural insights into C3 convertase activity of the classical pathway of complement"

+) current affiliation: Thermo Fisher Scientific, Electron microscopy division, Eindhoven, The Netherlands

### Supplementary Figures

Supplemental Figure 1:

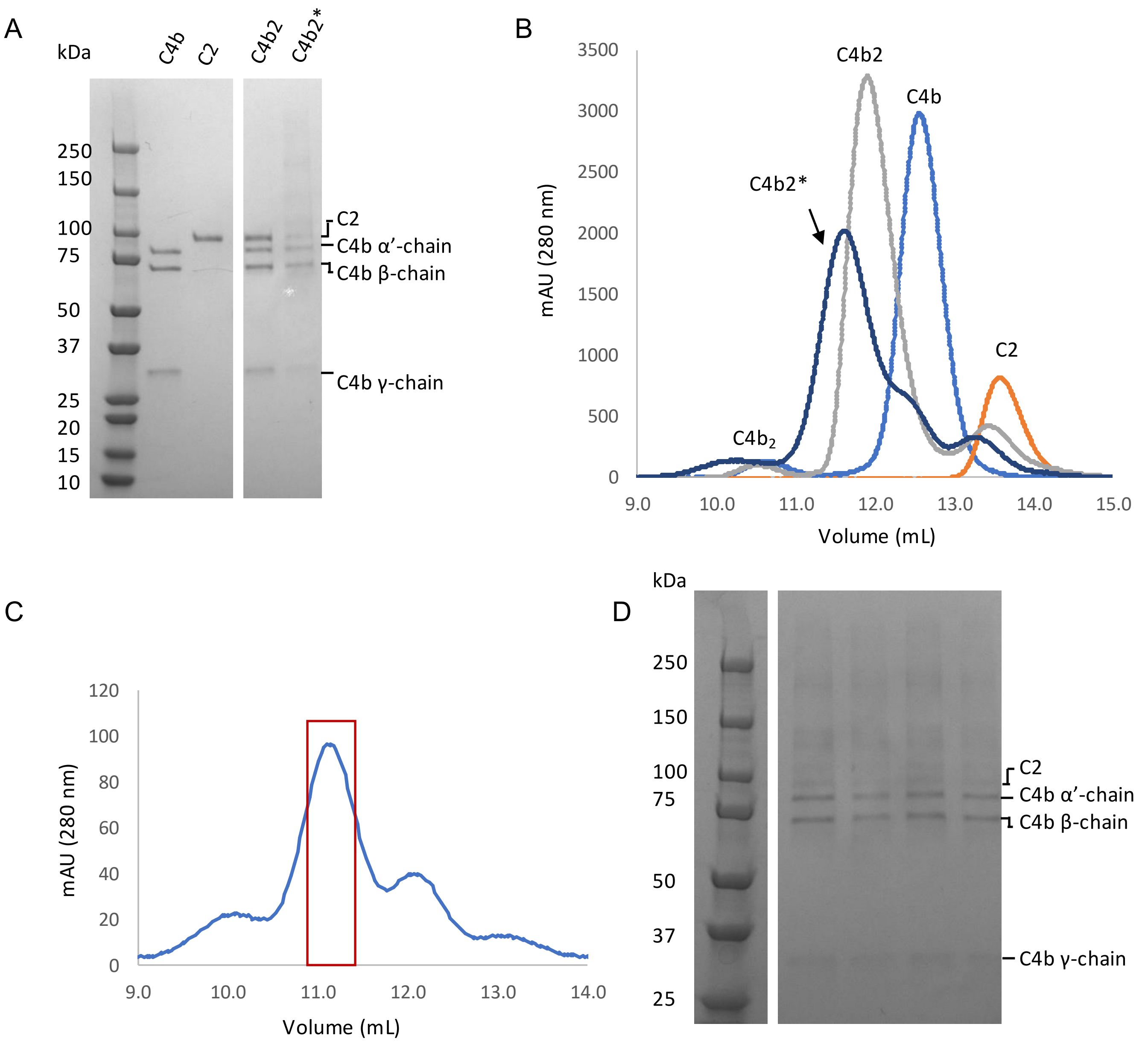


**Figure S1. Purification and cross-linking of proconvertase C4b2. (A)** SDS-PAGE gel and **(B)** analytic SEC of cross-linked C4b2 (C4b2*) with controls C4b, C2 and C4b2. **(C)** SEC of C4b2* with selected fractions for cryo-EM outlined in red and **(D)** SDS-PAGE gel of selected fractions.

Supplemental Figure 2:

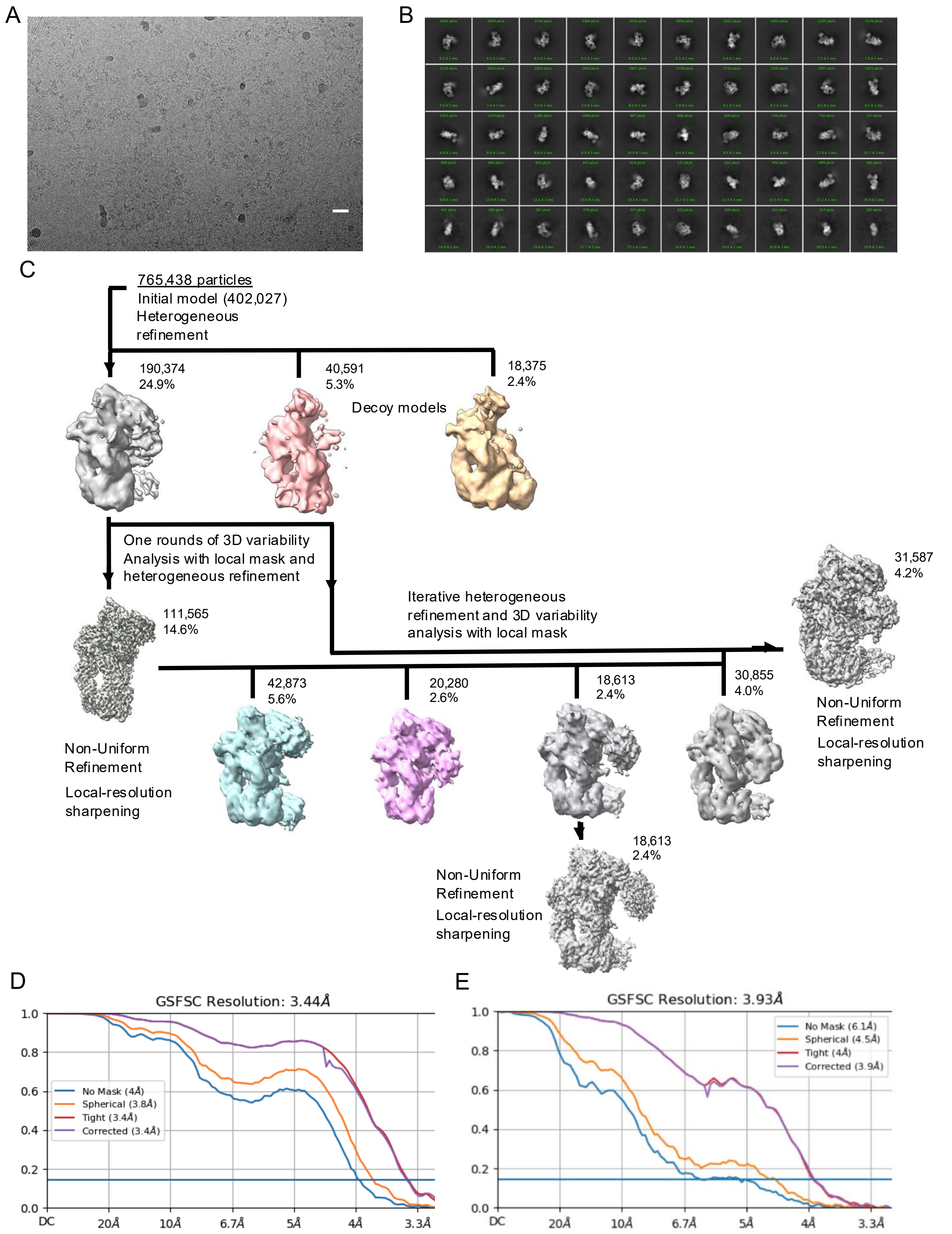


**Figure S2. Cryo-EM image processing of the proconvertase C4b2 dataset. (A)** Micrograph of cross-linked C4b2 particles in vitreous ice with a scale-bar length of 20 nm. **(B)** Selected 2D-class averages of the complex generated in cryoSPARC. **(C)** Cryo-EM processing workflow for C4b2 reconstructuin and refinement, including additional rounds of 3D variability analysis that reveals alternative SP domain conformations. **(D-E)** Fourier-shell correlation plots for the gold-standard (0.143) refined C4b2 reconstructions, computed from unmasked (blue), with a spherical mask (orange), tight mask (red), and corrected (purple) for C4b2 maps at 3.5 and 3.9-Å resolution, respectively.

Supplemental Figure 3:

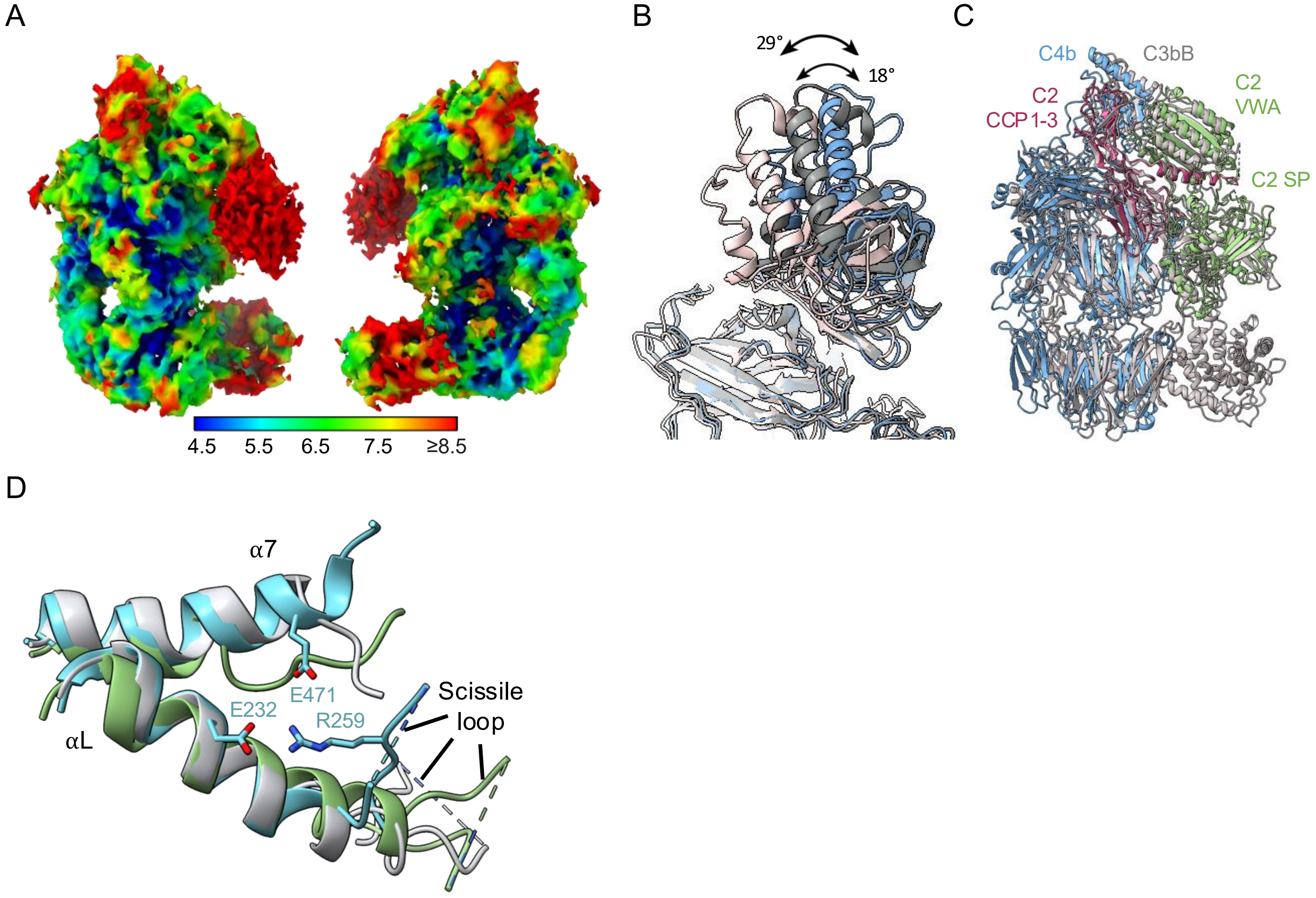


**Figure S3. Proconvertase C4b2. (A)** Cryo-EM density map of C4b2 at 4.2-Å resolution, reconstructed from a subset of particles selected via 3D variability analysis in CryoSPARC. SP domain appears partially unresolved and displaced. The map is coloured by local resolution as indicated by the bar below. **(B)** Superposition free C4b (grey, pdb 5JPN) and C4b from the C4b-hC4Nb8 nanobody complex (light pink, pdb 6YSQ), onto C4b of the C4b2 complex (blue). **(C)** Superposition of C3bB (grey, pdb 2XWJ) on C4b2 with C4b in blue, C2 CCP1-2 and αL in purple and C2 VWA-SP in green. **(D)** Superposition of VWA domains from C4b2 (green), C3bB (grey), and CVF-B (cyan, pdb 3HRZ), showing helices αL and α7. Scissile loops in C4b2 and C3bB are exposed and disordered and exposed, while in CVF-B the FB P1-residue Arg259 is buried and interacts with αL (Glu232) and α7 (Glu471).

Supplemental Figure 4:

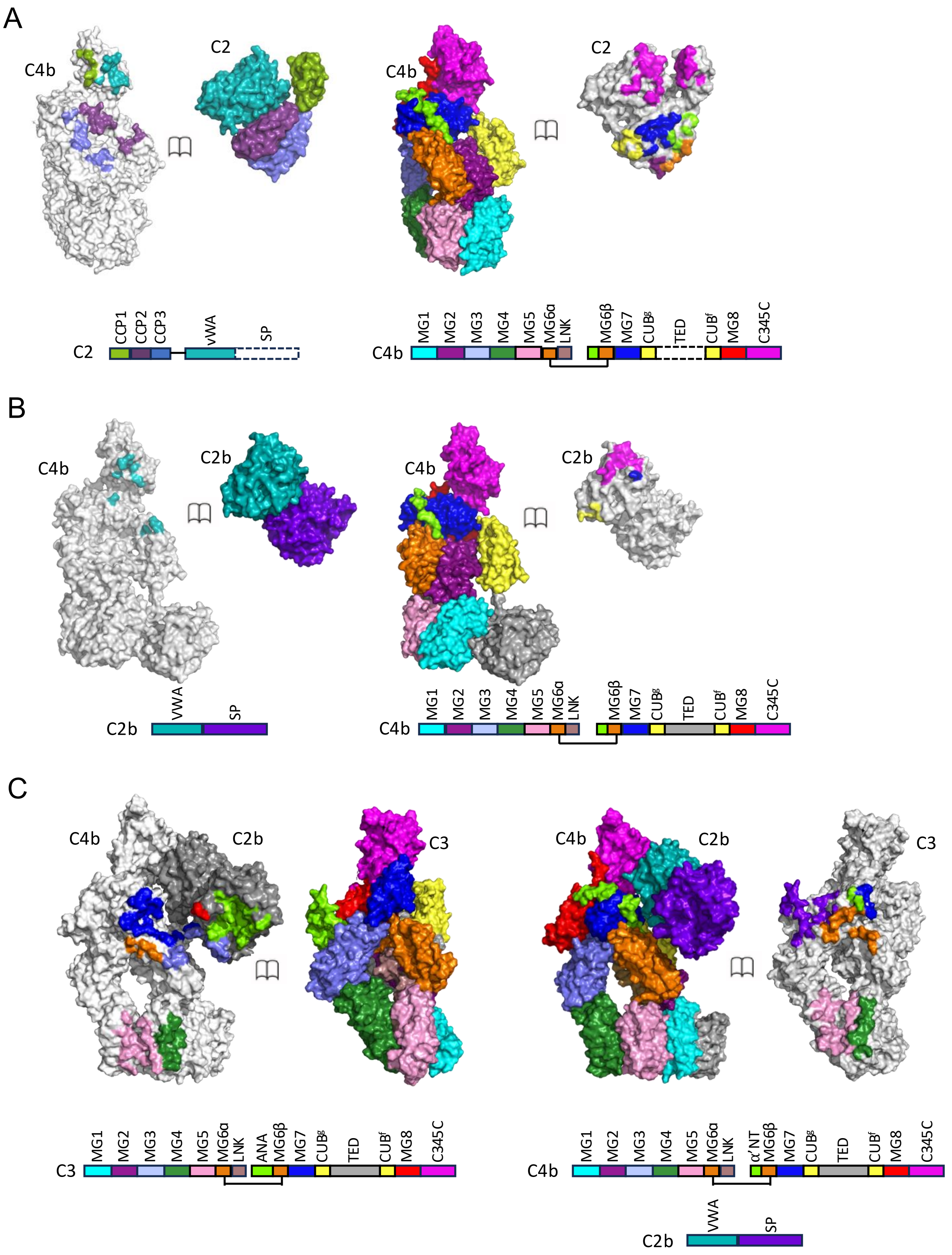


**Figure S4. Contact footprints of C4b2, C4b2b and C4b2bC3.** Opened book view of the contact footprints of **(A)** C4b2, **(B)** C4b2b and **(C)** C4b2b-C3. Domain colouring as indicated by the bar diagrams (domains missing in the model are indicated by dashed white bars).

Supplemental Figure 5:

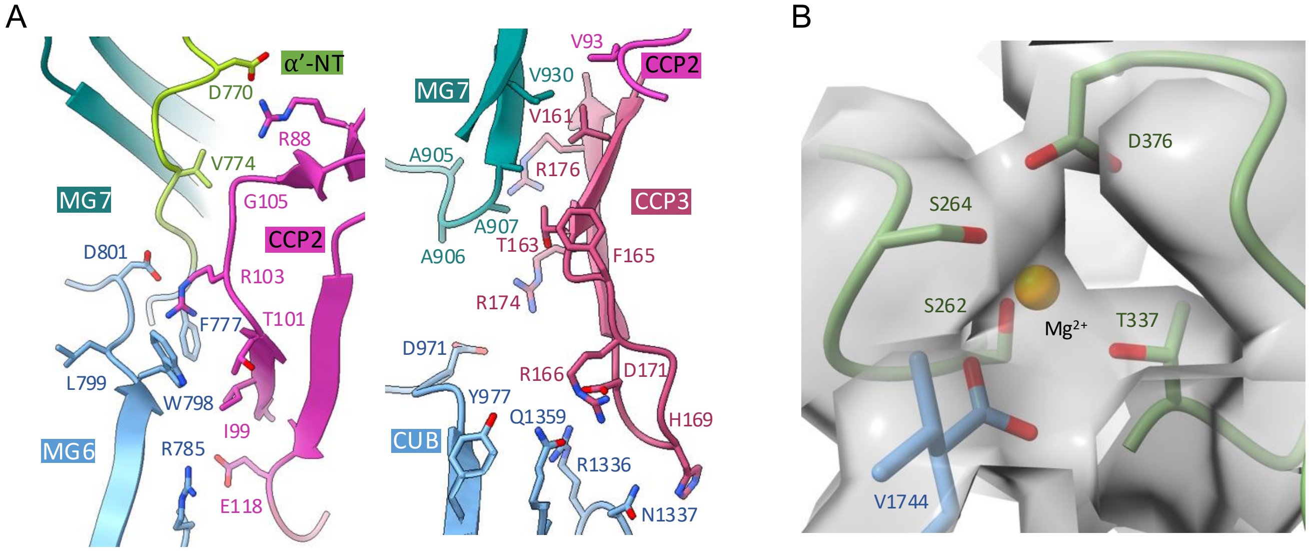


**Figure S5. Interactions between C4b and C2 in C4b2. (A)** Interactions between C4b MG6 (light blue), MG7 (blue-green) and α'NT (green) with C2 CCP2 (purple) (left panel) and C4b MG7 (blue-green) and CUB (blue) with C2 CCP2 (purple) and CCP3 (red) (right panel). **(B)** Cryo-EM density map shown in semi-transparent grey of the metal-ion dependent adhesion site (MIDAS) with C2 VWA interacting residues in green, C- terminal tail of C4b γ-chain in blue and magnesium ion as an orange sphere.

Supplemental Figure 6:

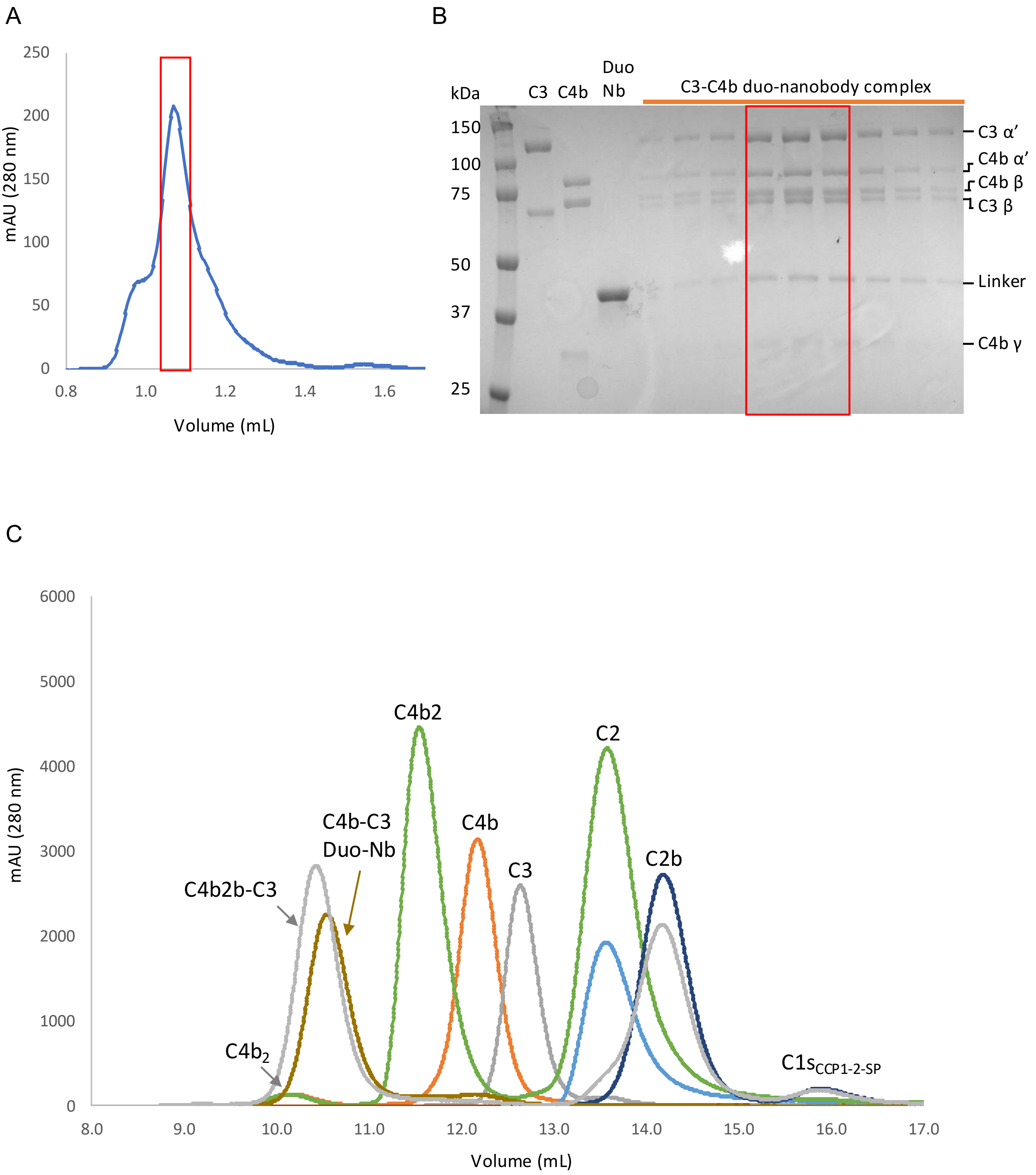


**Figure S6. Formation and purification of convertase-substrate complex C4b2bC3. (A)** SEC purification of anti-C4b and anti-C3 duo-nanobody complex with selected fractions outlined in red. **(B)** SDS-PAGE gel of selected fractions outlined in red and C4b, C3 and duo-nanobody linker as controls. **(C)** Analytical SEC of convertase-substrate C4b2b-C3 formation (grey) with cleaved C1s, C2, C2b, C3, C4b, C4b2 and duo-nanobody as controls. A cartoon image of the eluted protein or protein complex is shown on top of the elution peak to facilitate the visualization and interpretation of the chromatogram.

Supplemental Figure 7:

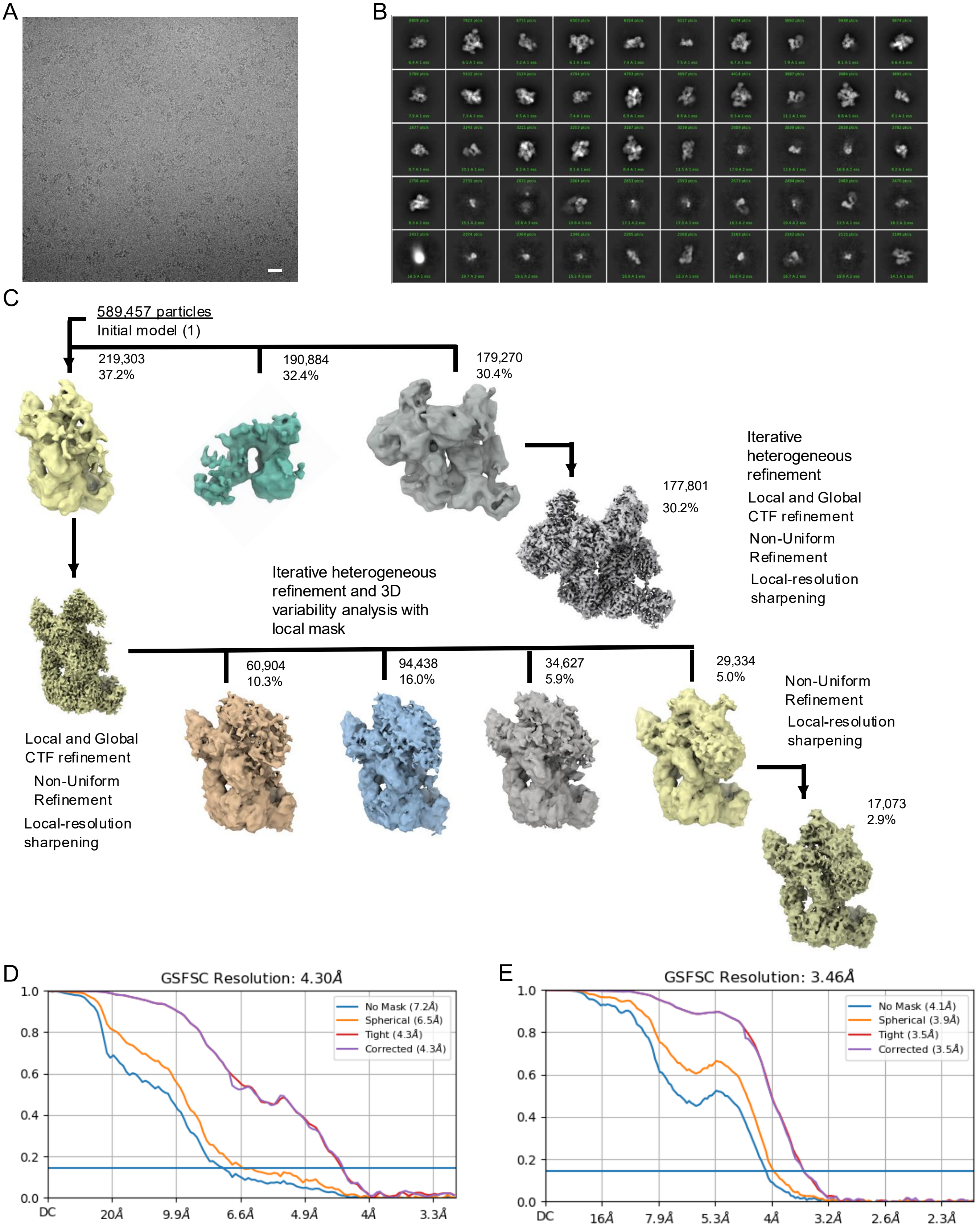


**Figure S7. Cryo-EM image processing of convertase C4b2b and convertase-substrate complex C4b2b-C3. (A)** Micrograph of C4b2b and C4b2b-C3 particles in vitreous ice with a scale bar length of 20 nm. **(B)** Selected 2D-class averages of the complexes generated in cryoSPARC. **(C)** Cryo-EM processing workflow for Proteins C4b2b and C4b2b-C3, including extended 3D variability analysis for protein complex C4b2b to resolve C2b domains. **(D-E)** Fourier-shell correlation plots for the gold-standard (0.143) refined C4b2b and C4b2bC3 reconstructions, respectively, computed from unmasked (blue), with a spherical mask (orange), tight mask (red), and corrected (purple).

Supplemental Figure 8:

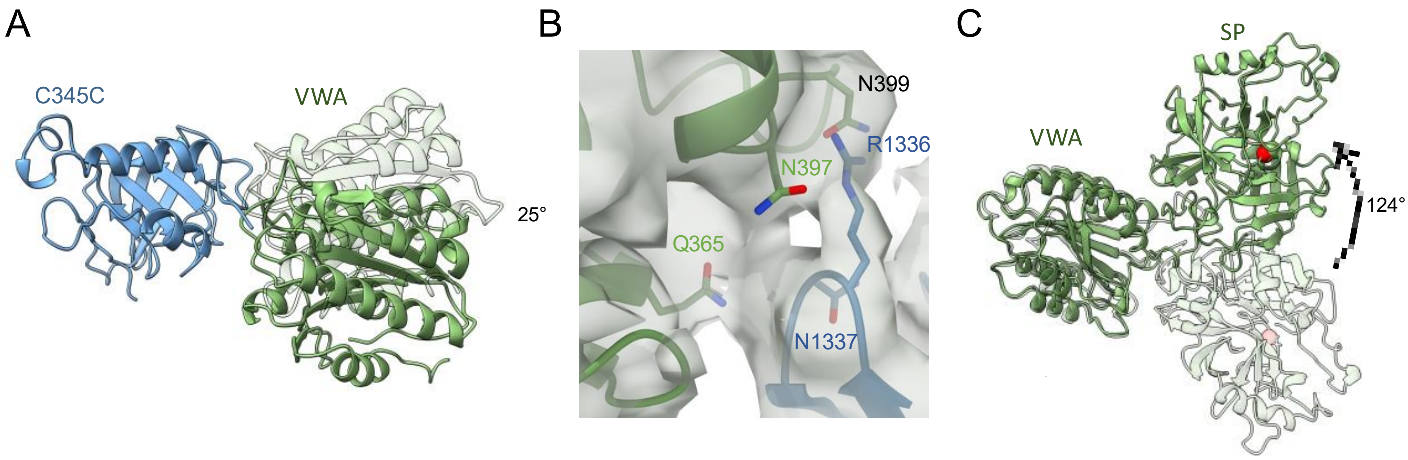


**Figure S8. Convertase, C4b2b. (A)** Superposition of C4b C345C domain (blue) of C4b2b and C4b2, with domain VWA shown in solid and transparent green, respectively, showing a 25° rotation. (**B**) Cryo-EM density map shown in semi-transparent grey; showing contact point of C4b CUB and C2b VWA. **(C)** Superposition of VWA domain of C4b2b and C4b2, with C2b VWA-SP shown in solid green and C2 VWA-SP in transparent green. SP domain rotates 124° repositioning and reorienting the enzyme active site. Cα of S679A is indicated by red spheres for both proteins.

Supplemental Figure 9:

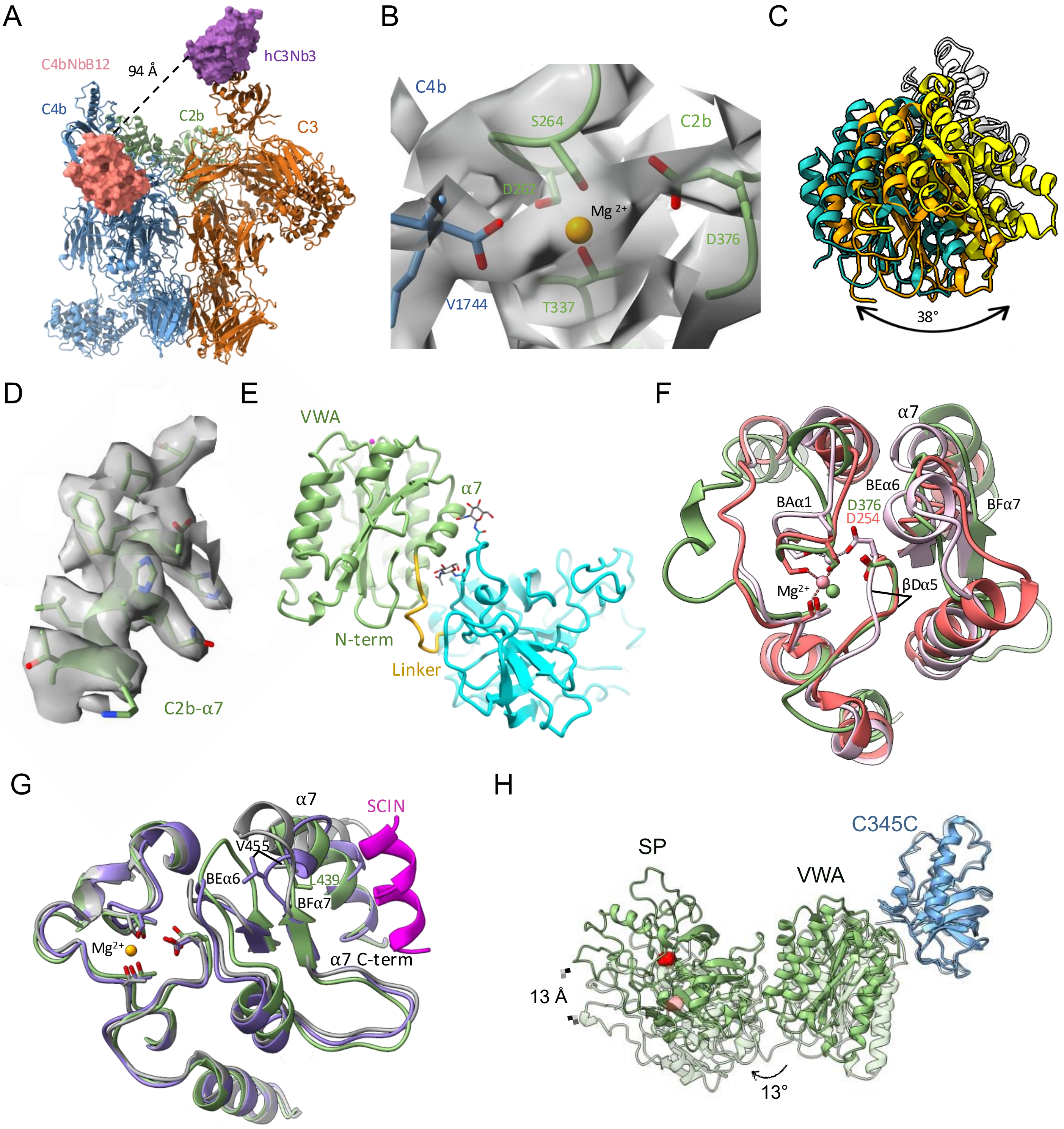

**Figure S9. Convertase-substrate C4b2b-C3. (A)** C4b2b-C3 showed in cartoon representation showing C4b (blue), C2b (green) and C3 (orange). Anti-C4b nanobody C4bB12 and anti C3 nanobody hC3Nb3 are shown in surface representation in pink and purple, respectively. The black dotted line indicates the minimum distance between the C and N-terminus of anti-C4b and anti-C3 duo-nanobody, resp. **(B)** Cryo-EM density map shown in semi-transparent grey of the MIDAS site with C2b residues shown in green, C-terminal tail of C4b in blue and $Mg^{2+}$ in orange. **(C)** VWA domains of C4b2 (yellow), C4b2b (orange) and C4b2b-C3 (light blue) after superposing their C4b C343C domains. **(D)** Cryo-EM density map shown in semi-transparent grey of the $\alpha$7 helix (green) of C2b in C4b2b-C3 complex. **(E)** C2b of C4b2b-C3 in cartoon representation with VWA (green), VWA-SP linker (orange) and SP (cyan) and modelled N-glycosylations (grey). **(F)** Superposition of integrin $\alpha$2 I domain, in its unbound (light pink, pdb 1AOX), and ligand-bound state (dark pink, pdb 1DZI) onto C2b VWA (green) of C4b2b-C3; corresponding $Mg^{2+}$ ions shown in respective colours. **(G)** FB VWA domains of FB from C3bB (grey, pdb 2XWJ) and Bb from $(C3bBb\text{-}SCIN)_2$ (purple, pdb 2WIN) with its interaction with SCIN (pink) superimposed onto C2b VWA (green) of C4b2b-C3 complex and its $Mg^{2+}$ (orange). **(H)** C2b VWA-SP in C4b2b-C3 and C4b2b in solid and transparent green, respectively, after superposing their C4b C345C. C2b rotates 13° from free to substrate-bound convertase yielding displacement of S679A C$\alpha$ by 13 Å.

Supplemental Figure 10:

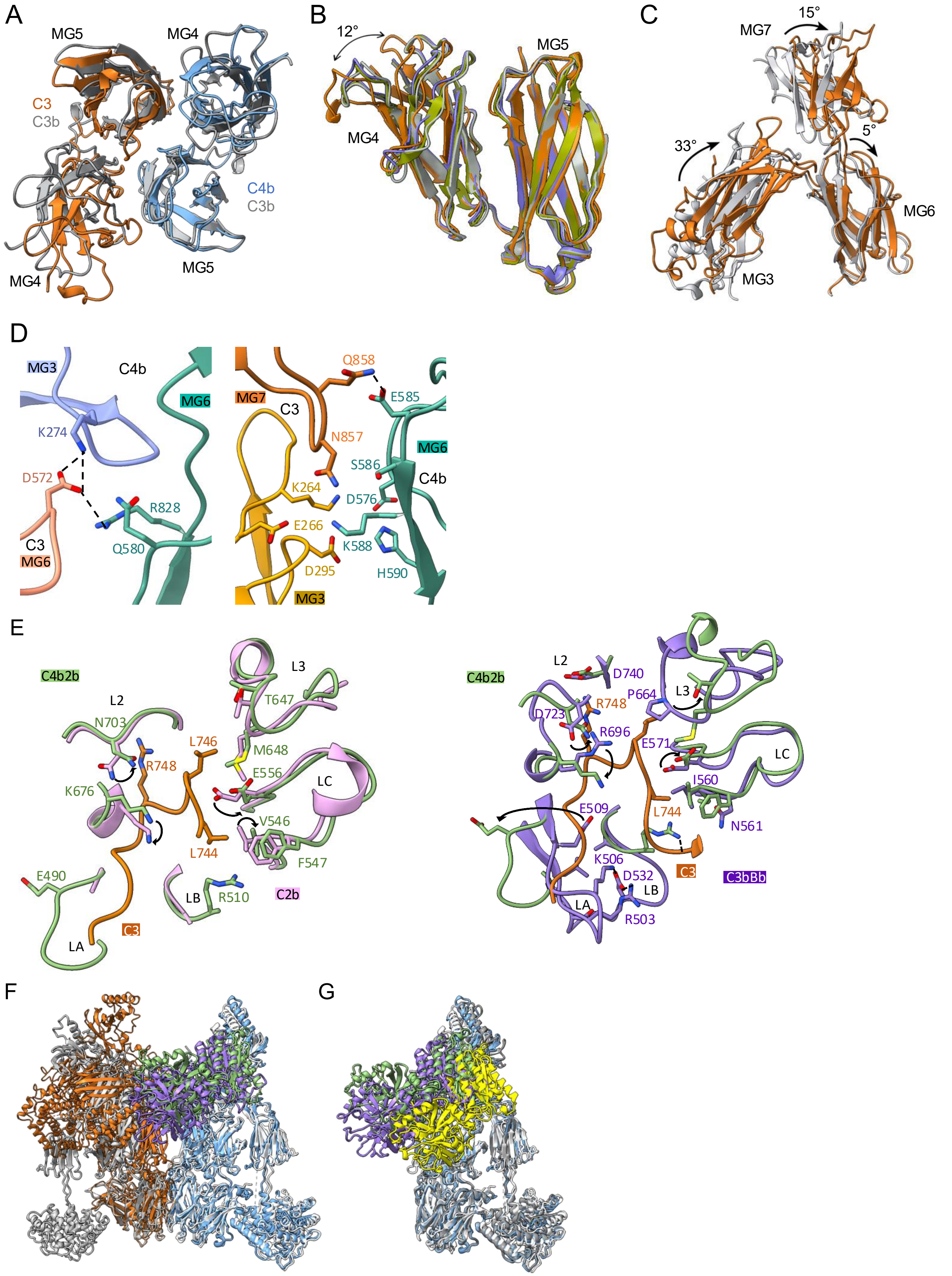

**Figure S10. Interactions between C3 and C4b2 in C4b2-C3**. **(A)** Cartoon representations of superposed C3b MG4-5 of $(C3bBb\text{-}SCIN)_2$ (grey, pdb 2WIN), C4b MG4-5 (blue) and C3 MG-5 (orange) of C4b2b-C3. **(B)** Superposition of MG4-5 of free C3 (grey, pdb 2A73), free C3b (green, 5FO7), C3b in $(C3bBb\text{-}SCIN)_2$ (purple) and C3 (orange) in C4b2b-C3. **(C)** C3 domains MG3, 6 and 7 of free C3 (grey) and C3 in C4b2-C3 (orange) after superposing on their MG5 domains. **(D)** Interactions between C4b MG3 (purple) and MG6 (teal) with C3b MG3 (yellow), MG6 (light pink) and MG7 (orange) in C4b2-C3. **(E)** Superposition of SP domain of free C2b (pink, pdb 2ODP) (left panel) and of Bb SP from $(C3bBb\text{-}SCIN)_2$ (purple, pdb 2WIN) (right panel) onto C2b (green) with bound C3-scissile loop (orange) of the C4b2b-C3 complex. **(F)** C4b2b-C3 (C4b in light blue, C2b in green and C3 in orange) and C3bBb-C3b (C3bBb, withC3b in white and Bb in purple, and C3b on substrate position in gray) taken from $(C3bBb\text{-}SCIN)_2$ (pdb 2WIN) superposed on the convertase C3b and C4b β-chains. **(G)** C3bBb of $(C3bBb\text{-}SCIN)_2$ (C3b in white and Bb in purple), free C4b2b (C4b in grey and C2b in yellow) and substrate-bound convertase C4b2b of C4b2b-C3 (C4b in light blue, C2b in green) superposed on their C3b or C4b β-chains.
